# Prediction of malaria mosquito species and population age structure using mid-infrared spectroscopy and supervised machine learning

**DOI:** 10.1101/414342

**Authors:** Mario González-Jiménez, Simon A. Babayan, Pegah Khazaeli, Margaret Doyle, Finlay Walton, Elliott Reedy, Thomas Glew, Mafalda Viana, Lisa Ranford-Cartwright, Abdoulaye Niang, Doreen J. Siria, Fredros O. Okumu, Abdoulaye Diabaté, Heather M. Ferguson, Francesco Baldini, Klaas Wynne

## Abstract

Despite the global efforts made in the fight against malaria, the disease is resurging. One of the main causes is the resistance that *Anopheles* mosquitoes, vectors of the disease, have developed to insecticides. *Anopheles* must survive for at least 12 days to possibly transmit malaria. Therefore, to evaluate and improve malaria vector control interventions, it is imperative to monitor and accurately estimate the age distribution of mosquito populations as well as total population sizes. However, estimating mosquito age is currently a slow, imprecise, and labour-intensive process that can only distinguish under-from over-four-day-old female mosquitoes. Here, we demonstrate a machine-learning based approach that utilizes mid-infrared spectra of mosquitoes to characterize simultaneously, and with unprecedented accuracy, both age and species identity of females of the malaria vectors *Anopheles gambiae* and *An. arabiensis* mosquitoes within their respective populations. The prediction of the age structures was statistically indistinguishable from true modelled distributions. The method has a negligible cost per mosquito, does not require highly trained personnel, is substantially faster than current techniques, and so can be easily applied in both laboratory and field settings. Our results show that, with larger mid-infrared spectroscopy data sets, this technique can be further improved and expanded to vectors of other diseases such as Zika and Dengue.

---

Between 2000 and 2015, insecticide-based control interventions targeting mosquito vectors averted an estimated 537 million malaria cases.^1^ Nevertheless, malaria still kills hundreds of thousands of people each year (445,000 in 2016), mainly in sub-Saharan Africa.^2^ Additionally, there is concern that progress on control may have stalled after more than a decade of success in global malaria control.^2^ Of major concern is the increase in insecticide resistance among mosquito populations throughout Africa,^3^ which is degrading the lethality and effectiveness of vector control tools, notably indoor residual spraying (IRS) and long-lasting insecticide treated nests (LLINs) which have been the cornerstones of malaria control in the past decades.^4^ Indeed, much of the effectiveness of LLINs and IRS comes from community-wide reductions in vector population size, not merely from preventing people from getting bitten.^5^

Measurement of female mosquito vector survival is the most important biological determinant of malaria transmission intensity.^6,7^ This is because malaria parasites *(Plasmodium* spp.) require more than 12 days of incubation inside the female mosquito vectors (extrinsic incubation period, EIP) before they become infectious.^8–11^ While there is uncertainty about mosquito survival in the field, crude estimates suggest the median lifespan of African malaria vectors is 7-10 days.^12^ Thus, only relatively old mosquitoes can transmit the parasite.^13^ As a result, even minor reductions in mosquito survival can have exponential impacts on pathogen transmission.^10,14^ Consequently, accurate and high-resolution estimation of both mosquito abundance and longevity is essential for the assessment of the impact of these and other control measures.

Despite the crucial importance of mosquito demography to vector control, there are few reliable tools for rapid, high-throughput monitoring of mosquito survival in the wild. Conventionally, mosquito age has been approximated by classifying females (the only sex that transmits malaria) into groups based on their reproductive status as assessed through observation of their ovarian tracheoles.^15^ This widely-employed technique distinguishes females who have not yet laid eggs (nulliparous) from those that have laid at least one egg batch (parous), with the latter group assumed to be older than the former because the gonotrophic cycle between blood feeding and oviposition takes ∼ 4 days. While useful for approximating general patterns in survival,^16^ this method is crude and cannot distinguish between females who have laid eggs only once or multiple times. Alternatively, more refined methods have been developed to estimate the number of gonotrophic cycles a female mosquito has gone through based on follicular relics or dilatations formed during each oviposition,^17^ although the conversion between gonotrophic cycles and actual age is imprecise (especially now that LLNIs are limiting regular access to blood-meals).^18^ While an improvement on the simple parity classification method, this approach is extremely technically demanding and time-consuming.^19^ Additionally, it is unsuitable for the analysis of sample sizes necessary for estimating mosquito population structure.^20^

Given these problems with ovary-based assessment, there has been significant investigation of alternative, molecular-based approaches to estimate mosquito age. These methods include: counting cuticle rings representing daily growth layers of the mosquito skeletal apodemes,^21^ chromatographic analysis of cuticular hydrocarbon chains,^22^ assessment of pteridines using fluorescence techniques,^23^ transcriptomic profiling,^24^ and mass spectrometric analysis of mosquito protein expression.^25^ However, thus far the level of accuracy, high cost, and/or need of highly trained users suggest that they might not be suitable for application in the field.

In addition to age, identification of mosquito species is crucial for estimation of malaria transmission dynamics. In Africa, the bulk of malaria transmission is carried out by members of the *Anopheles gambiae* sensu latu and *Anopheles funestus* groups.^26^ *An. gambiae* s.l. complex includes several cryptic species that can only be distinguished by molecular analysis.^27^-^29^ Despite being morphologically identical, members of this group vary significantly in behaviour, transmission potential, and response to vector control measurements.^30^ For example, two major vectors in the *An. gambiae* s.l. group, *An. arabiensis* and *An. gambiae,* differ in their propensity to enter and rest in houses, in host species choice, breeding conditions, resistance to insecticides, and tolerance to dry climates.^6,31,32^ Currently, *An. gambiae* s.l. species are best distinguished by polymerase chain reaction (PCR) methods,^33,34^ which are time-consuming and expensive, and can thus only be carried out on a subsample of mosquitoes collected during entomological surveillance. Alternative techniques have been developed such as isoenzyme electrophoresis^35^ or chromatography of cuticular components,^23^ but these are also very laborious and have weak discriminatory power.^36^

Non-PCR-based methods often rely on structural and chemical differences in the cuticle between species and age groups. In particular, near-infrared spectroscopy (NIRS) has been evaluated as a general strategy for the analysis of otherwise cryptic mosquito traits. The NIRS technique appears to represent a step forward in the study of mosquitoes since new methods were developed^37^ in an attempt to classify *Anopheles* mosquitoes as older or younger than 7 days,^38^ to discriminate between *An. gambiae* s.s and *An. arabiensis*,^38^ or to detect whether *Aedes aegypti* mosquitoes carry the Zika virus.^39^ The appeal of this approach is that it does not require reagents, and holds promise as a fast, practical and cost-effective method for entomological surveillance. Nevertheless, recent results have raised concerns that it may lack discriminatory power when the mosquitoes analysed have not been fed and reared homogeneously.^40^ Also, due to the impossibility of using a calibration dataset generated with one population of mosquitoes, the approach fails to predict properties of a different mosquito population.^41^

Here we propose that these limitations of NIRS can be overcome by shifting the range of analysis (4,000-25,000 cm^-1^) to another region of the infrared spectrum. Specifically, we propose the use of the mid-infrared region (400-4,000 cm^-1^) where the light interacts with the fundamental vibrations of the biomolecules present in the cuticle, creating an absorption spectrum of discrete well-delineated bands. Thus, the mid-infrared region provides a wealth of information not present in the near-infrared range, where only broadened combinations of the overtones of the fundamental vibrations appear.^42^ Mid-infrared spectroscopy (MIRS) has therefore several advantages for this application since it allows the capture of contributions of different biochemical components to the spectrum and their variations among mosquitoes with different attributes.^43,44^ However, since the mid-infrared spectral bands are affected in non-trivial ways by the development of a mosquito and the changing composition of the cuticle, it is not possible to predict traits by simply monitoring changes in band intensities.^45^

Here, we show that the use of supervised machine learning^46^ allows the determination of the age and species of two major malaria vectors, *An. arabiensis* and *An. gambiae*, with unprecedented speed and accuracy from the information contained in their mid-infrared spectra. This is possible because machine learning, unlike standard statistical approaches, can recognise the complex relationships in these traits (mosquito species and mosquito age) and disentangle them from other irrelevant variation.^47^-^49^ We demonstrate that as a result, we are able to reconstruct age distributions of mosquito populations with unprecedented reliability. The technique we propose here solves the drawbacks of the methods described previously, including being more time efficient (an analysis takes less than one minute per mosquito), less expensive, and requiring neither reagents nor highly trained operators, therefore constituting a key improvement in the tools for mosquito surveillance and evaluation of interventions.

## Results

A ‘field-friendly’ protocol to kill and store mosquitoes for infrared (IR) spectroscopy was established as described in Supplementary Note 1. In brief, laboratory-reared female *An. gambiae* and *An. arabiensis* mosquitoes of different ages and physiological states were killed by exposure to chloroform for 30 minutes. As chloroform evaporates and does not interact with the mosquito cuticle, the IR spectra were not affected by this chemical (Supplementary Figure 1). This method is more practical in the field than killing mosquitoes with CO2 or by freezing them at −20°C. Dead mosquitoes were then stored in 20 ml transport tubes with silica gel to dry them out. Removal of water from samples is essential, as it uncovers parts of the IR spectrum that would otherwise be hidden by the intense IR absorption of water (Supplementary Figure 2). Water IR absorption bands disappeared from *An. gambiae* and *An. arabiensis* mosquitoes after storage with silica gel at 4°C for one and two days, respectively. In addition, this drying method preserved the mosquitoes from decomposition for more than 10 days (Supplementary Figure 3). Alternative drying methods such as drying in an oven at 80°C was shown to affect IR spectra (Supplementary Figure 2) and therefore were not used.

The far-(30-400 cm^-1^), mid-(400-4,000 cm^-1^), and near-infrared (4,000-10,000) regions of the mosquito spectra were compared (Supplementary Figure 4). The far-and near-infrared regions were essentially featureless in dried mosquitoes, unlike the spectra observed in the literature^37,39,40,50,51^ that show the intense signals of liquid water as they were measured in undried mosquitoes (Supplementary Figure 5). However, the mid-infrared region showed a large number of well-defined intense peaks, which are easily identifiable as coming from the chemical components of the cuticle (Supplementary Table 1). Three different IR spectral sampling techniques were investigated: diffuse reflectance, transmission, and attenuated total internal reflection (ATR, see Supplementary Note 2). ATR spectroscopy produced the best-defined and most reproducible spectra in the mid-IR region (Supplementary Figure 6). ATR also allowed the measurement of different parts of the mosquito body (e.g., head or abdomen) that have slightly different IR spectra (Supplementary Figure 7). It also has superior signal-to-noise ratios allowing acquisition of the spectra in 45 seconds.

It is estimated that by using the ATR sampling technique in the mid-IR, the light penetrates the sample by 3-22 μm (see Supplementary Note 3). As the cuticle of a mosquito is approximately 2-5 μm thick,^21,52^ the measured spectra encompass the outer shell and part of the interior of the insects. As the cuticle is mainly composed of chitin, proteins, and lipids, the spectra of these substances individually were compared with the whole-mosquito spectra (Figure 1). This allowed the assignment of the main vibrational modes of the mosquito cuticular constituents to each element (Supplementary Table 1). As the cuticular chemical composition is known to change with species and age,^53,54^ the relative magnitudes of these vibrational bands are expected to change. To quantify this change, 17 wavenumbers in the IR spectrum were selected corresponding to 13 well-defined vibrational absorption peaks (contributed in different proportions by the three main constituents) and 4 troughs (that provide information on spectrum intensity and offset). These 17 wavenumbers were then used for training machine learning models (see below).

**Figure 1.**
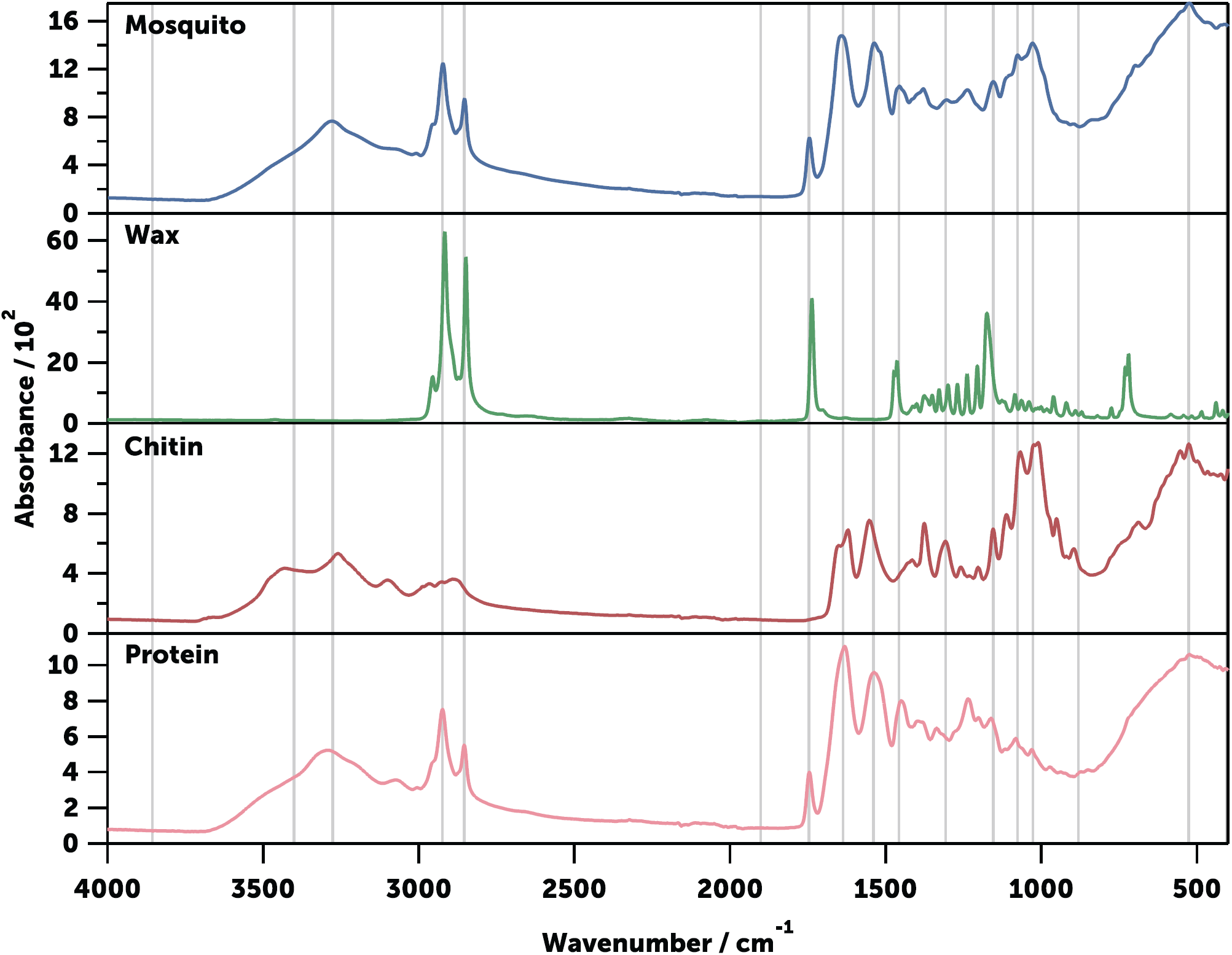
Typical mid-infrared spectrum of an Anopheles mosquito. Shown are An. gambiae (gravid, 9 days-old, blue) and its main chemical constituents wax (arachidyl dodecanoate, green), chitin (from shrimp shells, red), and protein (collagen from bovine Achilles tendon, pink). The wavenumbers selected for the machine learning are indicated with a grey line (Supplementary Table 1).

To develop a MIRS-based method to determine the age and species of *An. gambiae* and *An. arabiensis*, mosquitoes were reared under laboratory conditions (see Supplementary Note 1) and collected at ages ranging from 1 to 17. To model the variability typical of the wild, female mosquitoes in a range of physiological states were analysed including those that were had developed eggs (gravid), or had laid eggs but were not blood-fed (sugar fed). In most cases, over 40 mosquitoes per age and physiological condition from each species were analysed (Supplementary Table 2).

A total of 1,522 *An. gambiae* and 1,014 *An. arabiensis* spectra from different ages and conditions were used to train supervised machine-learning models (see Supplementary Note 4). Five algorithms were tested on the dataset to predict mosquito species (Supplementary Figure 8A). This initial approach identified logistic regression (LR) as the most accurate approach. We generated 100 bootstrapped models which, when aggregated (bagged), predicted the species identity of *An. gambiae* and *An. arabiensis* with 76.8 and 76.6% accuracy, respectively. To increase the accuracy of the prediction while retaining the stability and generalisability afforded by bagging, the 10 best models among them were selected, which achieved 82.6% accuracy (Figure 2). These results demonstrate that the MIRS signal is strongly indicative of mosquito species and can be used to distinguish between species in a more time and cost-efficient method, although currently with lower accuracy, than standard PCR methods.

**Figure 2.**
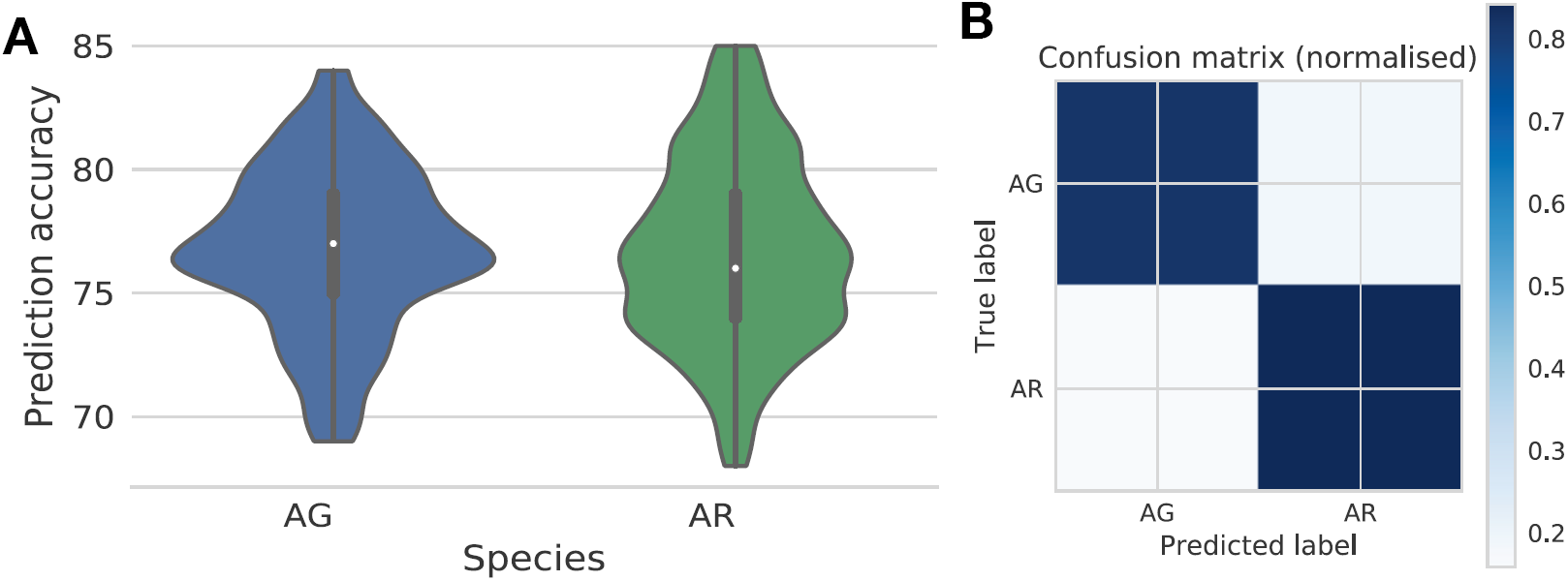
Prediction of mosquito species using mid-infrared spectra. (A) Distribution of per species prediction accuracies of 100 optimized models. (B) Confusion matrix showing the proportion of accurate (diagonal) classification of mosquitoes as either An. gambiae (AG) or An. arabiensis (AR) using the 10 best logistic regression models trained on repeated stratified random subset using 70% of all mosquitoes sampled and tested on the remaining 30% (n = 2,536).

After the development of the species-prediction model, a similar supervised machine-learning approach was used to model the chronological age for a given mosquito species. Mosquitoes were screened every second day after emerging as adults, and models trained on the same set of 17 wave-numbers as above. It was found that LR again performed best for both species in correctly mapping wavenumber intensities to mosquito age (Supplementary Figure 8B and C). To train, optimize, and validate the models, the full dataset was partitioned into an age-structured validation set retained for later use in population models (see below), and the remaining samples were then randomly split into stratified 70%/30% training and test sets for model tuning. The accuracy in predicting each chronological age varied over the mosquito lifespan and between species, ranging from an average of 15% to 97% for *An. gambiae* and 10% to 100% for *An. arabiensis* (Figure 3). It was found that the chronological age of young and old mosquitoes was generally more accurately predicted than intermediate ages, although there were some differences between species. These results suggest that the MIRS-based approach developed here can predict the chronological age of each species from 1 to 15 days old, as well as providing the confidence of prediction for each age class. Furthermore, a trade-off was observed between the granularity of the prediction and its accuracy: models trained on daily scans (not shown) performed worse than if we allowed the mosquitoes to age 2 or 3 days between each scan, suggesting that the ageing of the mosquito cuticle varies between individuals and that the features used for training the models overlap between consecutive age classes (Supplementary Figure 9).

**Figure 3.**
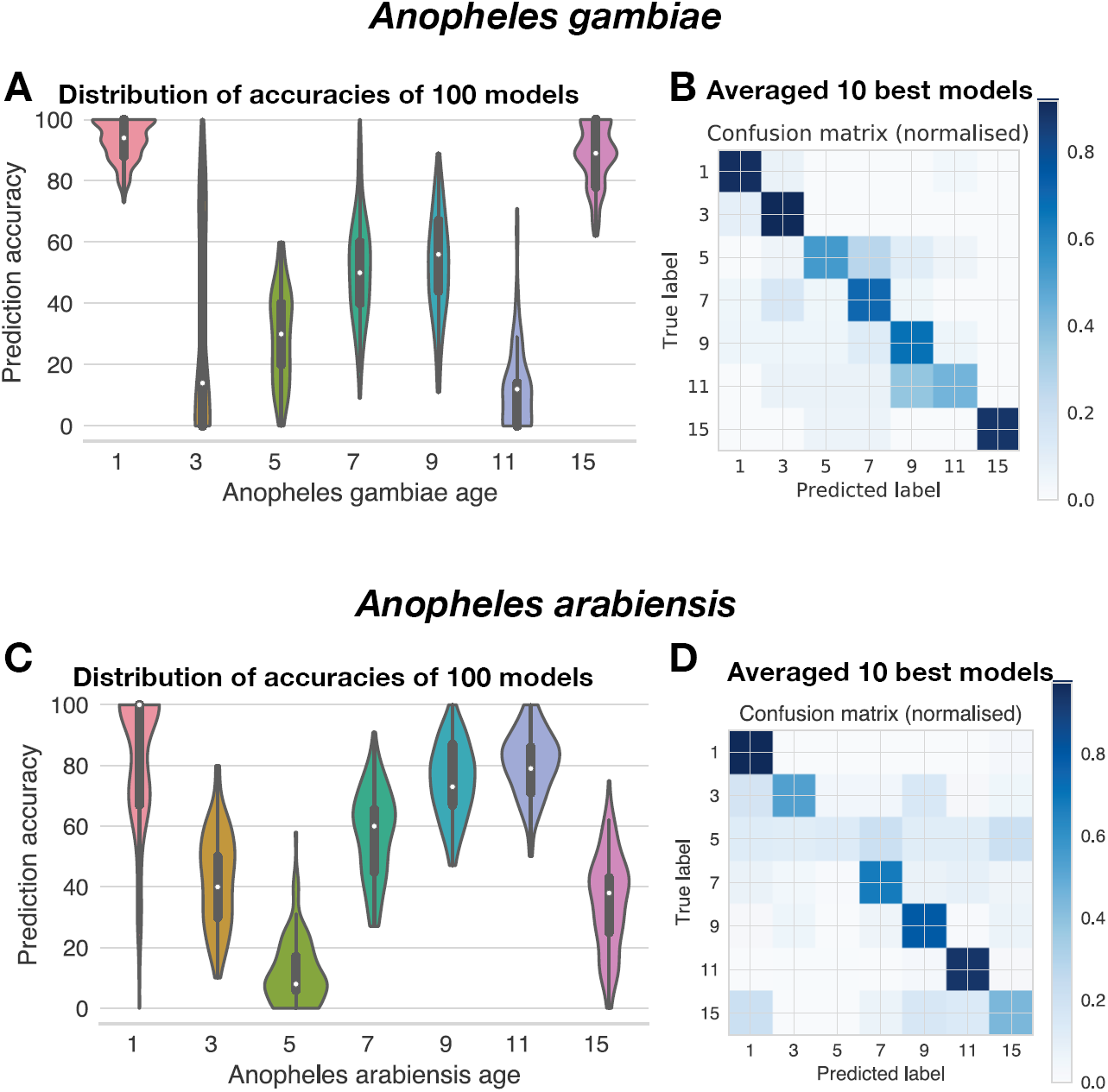
Prediction of An. gambiae (A-B) and An. arabiensis (C-D) age class using mid-infrared spectra. (A, C) Distribution of per age class prediction accuracies of 100 optimized models. (B, D) Confusion matrices showing the proportion of accurate (diagonal) classification of mosquitoes as either 1, 3, 5, 7, 9, 11, or 15 days old using the 10 best logistic regression models trained on repeated stratified random subsets using 70% of all mosquitoes sampled, and tested on the remaining 30% (n = 681 for An. gambiae and n = 737 for An. arabiensis).

To monitor the efficacy of vector control interventions in the field, accurately describing the age distribution *(i.e.,* the summary demographic age structure of the local vector species population) is more important than knowing the age of any individual mosquito.^41^ Consequently, we tested how well the above age prediction models developed for *An. gambiae* and *An. arabiensis* could reconstruct the known age distribution of mosquito populations. Mosquito populations reflecting anticipated changes in mortality were used under two scenarios: natural mortality and increased mortality due to vector control interventions.

Consistent with natural mosquito populations, but unlike our training dataset, field sampling would not produce age-balanced sample sizes, but rather very unbalanced age groups. Furthermore, it would be highly desirable to use our models to measure the impact of vector control interventions on mos-quito-population age structures. However, because no real datasets of a true mosquito population age structure exist, the age structures of *An. gambiae* and *An. arabiensis* were modelled based on their reported average daily mortality^16,55^ and assuming an intervention that increased the mortality of adult females four-fold after a first blood meal (∼3 days after adult emergence).

In these simulations, a starting population of 1,000 female mosquitoes was used, with the population at each subsequent day being calculated as a proportion of the previous day, with survival rates for each species estimated from reports on field studies (Figure 4A,D).^16,55^ The resulting age-structured populations were then used to randomly sample replicates in the corresponding proportion for each age class used in our MIRS-based prediction models (Figure 4B-F, grey bars; n = 122 for *An. gambiae* and n = 42 for *An. arabiensis).* The models trained above were then used to predict age classes from the MIRS of this age-structured population (Figure 4B-F, orange bars).

**Figure 4.**
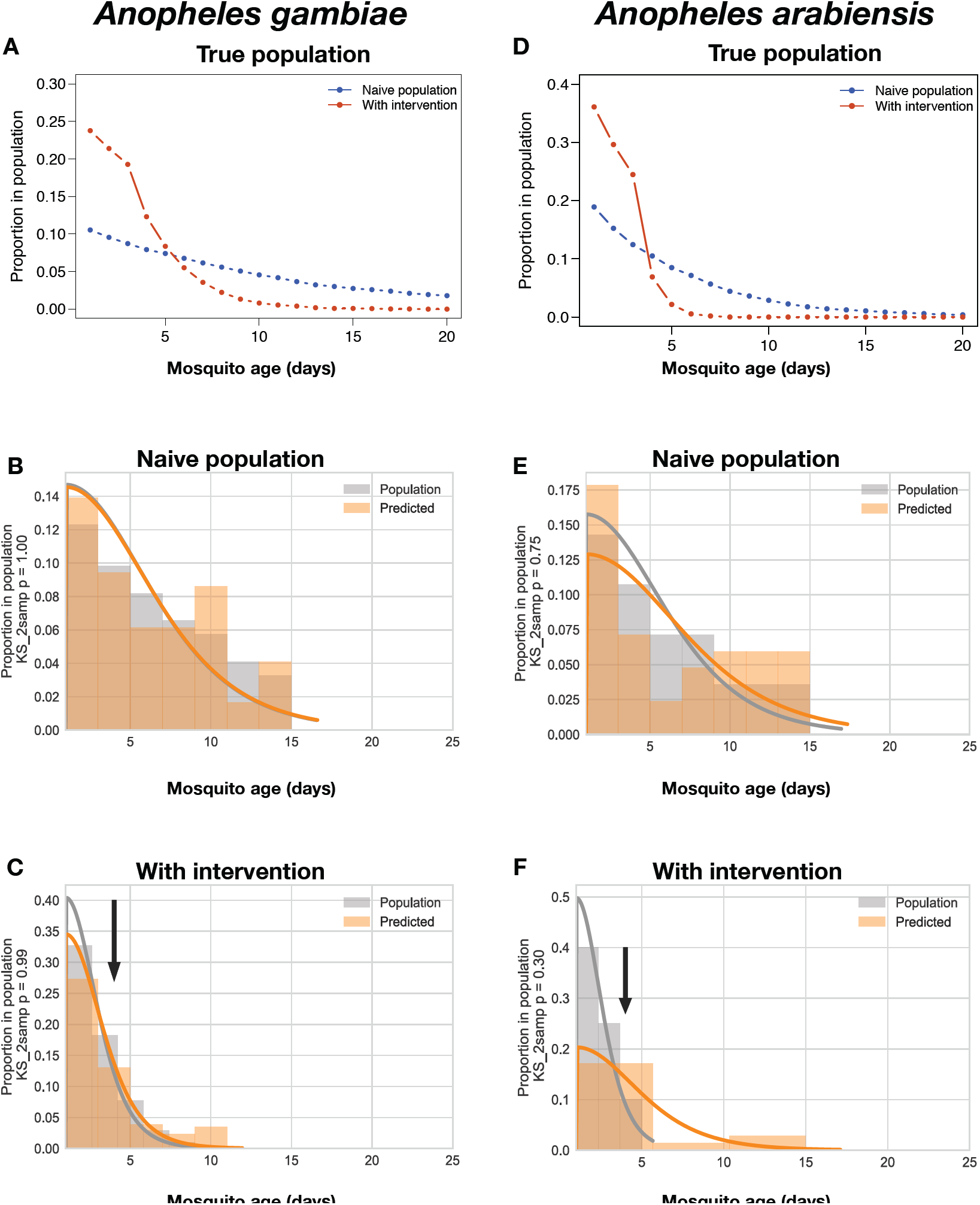
Reconstruction of the age structure of simulated populations of An. gambiae and An. arabiensis mosquitoes sampled from simulated pre-and post-treatment populations. Population age structures of An. gambiae (A-C) and An. arabiensis (D-F) were generated using an age structure population model assuming survival rates of 0.91 (An. gambiae, A) or 0.82 (An. arabiensis, D), under two common scenarios: naive untreated populations (blue lines), and populations in which a simulated vector control program resulted in 4x daily mortality of mosquitoes after day 3 (see Supplementary Note 5 for details). The proportions of each age class were extracted from those simulated populations (A, D), and used to build datasets that are representative of a field-sampled population survey (grey bars in B, C, E, and F). The resulting age-structured dataset **was** then used as the test set for our age-predicting machine learning models (see Figure 3) and compared with the predicted age structure generated from those models (orange bars in B, C, E, and F). Finally, we fit a continuous probability distribution to the true (grey curve) and predicted (orange curve) for better generalization of our discrete model predictions to an exponentially decreasing age structure. Population distributions were compared using a 2-sample Kolmogorov-Smirnov test (KS_2samp), reported in the y-axis labels. A - Relative proportion of each age class in a simulated population of An. gambiae. B - Estimation of age structure of simulated population from (A) using best models from Figure 3B for An. gambiae (n = 130). C - Estimation of age structure of simulated population post-intervention from (A) using best models from Figure 3B for An. gambiae (n = 122). D - Relative proportion of each age class in a simulated population of An. arabiensis. E - Estimation of age structure of simulated population from (A) using best models from Figure 3D for An. arabiensis (n = 42). F - Estimation of age structure of simulated population postintervention from (A) using best models from Figure 3D for An. arabiensis (n = 45).

To test the ability of those models to reconstruct the age structure of the true population from our predicted age class frequencies, the age structures of the predicted (Figure 4B-F, orange bars) and true sampled populations (Figure 4B-F, grey bars) were modelled with the best fit half-logistic distribution for each species (grey and orange curves in Figs. 4 B,C and E,F; see also Supplementary Note 5). The true and predicted age distributions were statistically indistinguishable (Kolmogorov-Smirnov 2-sample test (KS test), p = 1 and p = 0.99 for *An. gambiae* pre-and post-intervention, respectively; p = 0.75 and p = 0.30 for *An. arabiensis* pre-and post-intervention, respectively). This approach shows that the algorithm can reconstruct the age structure with good accuracy. Furthermore, our models detected a shift in mosquito age structure consistent with the simulated impacts of the interventions (sampled from true population: KS test p < 0.0001 for *An. gambiae* and p = 0.004 for *An. arabiensis;* predicted population: p < 0.0001 for *An. gambiae* and p = 0.1 for *An. arabiensis*), suggesting that this MIRS-based approach holds great promise for robust measurement and estimation of the age structure of mosquito vector populations.

## Discussion

We developed an easy, inexpensive, and rapid method to efficiently determine the age and species of large numbers of *An. gambiae* s.l. mosquitoes *(An. gambiae* s.s. and *An. arabiensis*). Based on the supervised machine-learning analysis of their mid-infrared spectra, this method allows the monitoring of the mosquito species distribution and the determination of mosquito survival and its associated vectorial capacity, two crucial tasks critical to implement and assess malaria control strategies. This approach will strongly benefit the vector-control community because, unlike the current most widely used technique based on dissection, it can accurately determine the whole-age distribution of a mosquito population from the day of emergence until two weeks of age. In addition, this approach provides an accurate estimate on a finer scale of the proportion of mosquitoes within the older age classes responsible for transmission.

The use of the mid-infrared spectral region provides clear advantages over techniques using near-infrared. Foremost, it is possible to independently quantify the amount of different biochemical components as their vibrational bands appear at different wavenumbers. Furthermore, the spectra are measured faster and the MIRS bands are more intense and have much greater definition. In contrast, the near-infrared spectrum of a mosquito is composed of few weak signals (Supplementary Figure 4) that are typically dominated by the much stronger vibrational overtone and combination bands of water (Supplementary Figure 5). As a consequence, analysis of the NIR spectrum of undried mosquitoes is unreliable, as the water signal is more dependent on the mosquito physiological state and environmental conditions than on other mosquito traits, such as species and age. It is therefore likely that the origin of the high accuracies claimed by NIR-based studies is due to the overfitting of the data. Indeed, recently reported analyses of NIRS data from different studies showed that the calibration and therefore predictive power could not be transferred between different datasets.^41^

We have shown that the variation of MIR spectra over mosquito age can be exploited by a machine-learning algorithm to predict the chronological age with a high degree of accuracy, and ultimately reconstruct population age structures of two important malaria vector species under simulated conditions of changing mortality risk due to vector control. Our algorithms accurately reconstructed age structures of both *An. arabiensis* and *An. gambiae,* in ways that were statistically indistinguishable from reality, and also detected shifts in mosquito age structure consistent with simulated impacts of interventions. To our knowledge, this is the first time that a proposed technique has had the ability to predict the age structure of a population or reconstruct such demographic patterns given a specific vector control intervention. The observed ability of MIRS to predict the change in mosquito death rate suggests that this approach will constitute an efficient tool for monitoring the efficacy of vector control interventions against mosquitoes. Future work will include larger datasets used for training in supervised machine learning, comprising field samples with different ecological conditions. The ecological variability of field samples has strongly limited the use of NIRS for age prediction.^40^ While the accuracy of MIRS-based approaches may also decline when moving from laboratory-reared to field mosquitoes, we predict that our method will be more robust due to the much greater information content and signal clarity that is obtained in spectra from MIRS relative to NIRS. In addition, it is notable that age was predicted accurately at the level of calendar day, for most age classes. We expect that varied mosquito responses to different field conditions, will result in more tractable physiological changes, further improving accuracies of our day-to-day age predictions relative to the laboratory samples. Besides, increasing the size and variability of the training set—which will permit using alternative machine learning techniques such as neural networks^56^—will facilitate the use of this MIRS-based approach for even higher throughput, accurate determination of mosquito age structure in vector surveillance programmes.

We have shown that MIRS can discriminate between morphologically identical *An. gambiae* s.l. species with ∼83% accuracy. Differences in behaviour and ecology among *Anopheles* species will determine the impact of specific vector control interventions.^6^ Currently the only way to distinguish morphologically identical *An. gambiae* s.l. species is by PCR,^33,34^ which requires trained staff, a well-equipped laboratory, is time-consuming, and expensive. The use of this MIRS-based approach will allow the determination of *An. gambiae* s.l. complex species without any molecular identification. While the observed accuracy of MIRS species prediction is still not comparable to the PCR precision, further work including a larger training set and field samples can increase the overall accuracy of this approach. In addition, the inclusion of other species of the *An. gambiae* s.l. complex will be necessary to implement this technique for field application. Nevertheless, these laboratory-based results, which included mosquitoes from different ages, physiological conditions, and cohorts, suggest that despite the ecological and life-history traits variation, MIR spectra contain a species-specific signature that the machine-learning algorithm can detect. Indeed, mass-spectrometry studies have shown that different species in the *An. gambiae* s.l. complex have quantitative differences in the cuticular hydrocarbon composition of their cuticle,^54^ which will affect the MIR spectra.

The biochemical signature obtained by MIRS from the mosquito cuticle provided information on both the species and the age. It may therefore be possible to obtain further information on other mosquito traits that alter the cuticular composition. Recently, a new insecticide resistance mechanism has been discovered in *An. gambiae*, which relies on an increased cuticle thickness that in turn reduces insecticide uptake.^57^ While this mechanism has been detected by electron microscopy, there are no other methods to measure this new trait, which could have profound epidemiological consequences. In the future, MIRS calibrations including cuticular resistant mosquitoes might be able to produce models to discriminate this insecticide resistant trait. In addition, infection with the *Plasmodium* malaria parasite might be detected by MIRS. Indeed, microorganism infections alter mosquito physiology and could directly or indirectly modify its cuticular composition. For example, in the dengue and Zika vector *Ae-des aegypti* mosquitoes, infrared spectroscopy has recently been developed to detect Zika virus,^39^ and the bacterial endo-7. symbiont *Wolbachia*.^43,50^

The accuracy and generalisability of the MIRS approach presented here to determine key mosquito traits shows that this tool can be used to assist the evaluation of vector control intervention in large mosquito populations. The inclusion of new species, larger sample sizes and field samples with variable ecological conditions is a prerequisite for the application ^9^ of this technique. It is worth noting that the cost of a portable FTIR MIR spectrometer is ∼$20-25,000, which is in the range of quantitative PCR machines used for species determination and/or insecticide resistance monitoring. However, in contrast to PCR analysis, no additional, ongoing costs for reagents and running costs are required once the core equipment is installed. Thus, this approach could be particularly valuable in resource limited settings.

The MIRS method presented here provides rapid and ac-12 curate information on *Anopheles* species (82.6%) and most importantly the mosquito age distribution. This is the first time that it has been possible to measure age distributions— including the effects of intervention—albeit with a resolution 13. of only two days. However, these results were obtained by training machine learning models with a relatively modest 14. number of mosquitoes (2,536). In future work, by employing multiple malaria research labs in Europe, Africa, and else-15. where, it will be possible to generate much larger MIRS datasets that can be used to train more sophisticated predictive models. This may then allow one to take advantage of deep learning methods with potentially further improved accuracy. The technique was applied here to malaria vectors but could clearly be expanded to other vector-borne diseases such as Zika, dengue, Lyme disease, leishmaniasis, or filariasis.

## Data availability

The mosquito MIRS data that support the findings of this study are available in Enlighten: Research Data Repository (University of Glasgow) with the identifier: http://dx.doi.org/10.5525/@@@

## Acknowledgements

This work was supported by EPSRC grants EP/J009733/1, EP/K034995/1, EP/N508792/1, and EP/N007417/1, and MRC grant MR/P025501/1. FO was also supported by a Wellcome Trust Intermediate Fellowship in Public Health and Tropical Medicine (Grant Number: WT102350/Z/13) and FB by an AXA RF fellowship (14-AXA-PD0C-130) and an EMBO LT fellowship (43-2014). We would like to thank Dorothy Armstrong and Elizabeth Peat for assistance with mosquito rearing and maintenance.

## Author contributions

MGJ, SAB, HMF, FO, FB, and KW wrote the paper, with contributions from PK, MD, FW, ER, TG, MV, LRC, AN, DJS, and AD. Spectral measurements were carried out by MGJ, PK, MD, FW, ER, and TG. The mosquitoes were grown by FB. Spectroscopy, wavenumber, associated data collection, and preliminary data analysis were carried out by MGJ, age models by MV, and statistics and machine learning by SAB.

## Additional information

Supplementary Information accompanies this paper at https://doi.org/10.1038/@@@@

## Competing financial interests

The authors declare no competing interests.

## References

1 Bhatt, S. et al. The effect of malaria control on Plasmodium falciparum in Africa between 2000 and 2015. Nature 526, 207–211 (2015).

2 WHO. World Malaria Report. (2017).

3 Hemingway, J. et al. Averting a malaria disaster: Will 20 insecticide resistance derail malaria control? Lancet 387, 1785–1788 (2016).

4 Protopopoff, N. et al. Effectiveness of a long-lasting 21 piperonyl butoxide-treated insecticidal net and indoor residual spray interventions, separately and together, against malaria transmitted by pyrethroid-resistant mosquitoes: a cluster, randomised controlled, two-by-22 two fact. Lancet 391, 1577–1588 (2018).

5 Hawley, W. A. et al. Community-wide effects of per-methrin-treated bednets on child mortality and malaria morbidity in western Kenya. Am J Trop Med Hyg 23 68, 121–127 (2003).

6 Pates, H & Curtis, C. Mosquito Behavior and Vector Control. Annu. Rev. Entomol. 50, 53–70 (2005).

7 Viana, M., Hughes, A., Matthiopoulos, J., Ranson, H. & Ferguson, H. M. Delayed mortality effects cut the malaria transmission potential of insecticide-resistant mosquitoes. Proc. Natl. Acad. Sci. 113, 8975–8980 (2016).

8 Beier, J. C. Malaria Parasite Development in Mosquitoes. Annu. Rev. Entomol. 43, 519–543 (1998).

9 Ohm, J. R. et al. Rethinking the extrinsic incubation period of malaria parasites. Parasites and Vectors 11, 1–9 (2018).

10 Smith, D. L. & McKenzie, F. E. Statics and dynamics of malaria infection in Anopheles mosquitoes. Malar. J. 3, 13 (2004).

11 Brady, O. J. et al. Vectorial capacity and vector control: Reconsidering sensitivity to parameters for malaria elimination. Trans. R. Soc. Trop. Med. Hyg. 110, 107–117 (2016).

12 Gillies, M. T. & Wilkes, T. J. A study of the age composition of populations of Anopheles gambiae Giles and Anopheles funestus Giles in north-eastern Tanzania. Bull. Entomol. Res. 56, 237–262 (1965).

13 Macdonald, G. Epidemiological basis of malaria control. Bull. World Health Organ. 15, 613–626 (1956).

14 Macdonald, G. The Epidemiology and Control of Malaria. (Oxford University Press, 1957).

15 Detinova, T. S. Age-grouping methods in diptera of medical importance, with special reference to some vectors of malaria. (1962).

16 Charlwood, J. D. et al. Survival and infection probabilities of anthropophagic anophelines from an area of high prevalence of Plasmodium falciparum in humans. Bull. Entomol. Res. 87, 445–453 (1997).

17 Polovodova, V. P. Age changes in ovaries of Anopheles and methods of determination of age composition in mosquito populations. Med. Parazitol. i Parazit. Bolezn. 10, 387–395 (1941).

18 Yakob, L. & Yan, G. Modeling the effects of integrating larval habitat source reduction and insecticide treated nets for malaria control. PLoS One 4, e6921 (2009).

19 Anagonou, R. et al. Application of Polovodova’s method for the determination of physiological age and relationship between the level of parity and infectivity of Plasmodium falciparum in Anopheles gambiae s.s, south-eastern Benin. Parasites and Vectors 8, 1–9 (2015).

20 Hoc, T. Q. & Charlwood, J. D. Age determination of *Aedes cantans* using the ovarian oil injection technique. Med. Vet. Entomol. 4, 227–33 (1990).

21 Schlein, Y. & Gratz, N. G. Determination of the age of some anopheline mosquitos by daily growth layers of skeletal apodemes. Bull. World Health Organ. 49, 371–375 (1973).

22 Gerade, B. B. et al. Field Validation of Aedes aegypti (Diptera: Culicidae) Age Estimation by Analysis of Cuticular Hydrocarbons. J. Med. Entomol. 41, 231–238 (2004).

23 Wu, D. & Lehane, M. J. Pteridine fluorescence for age determination of Anopheles mosquitoes. Med. Vet. Entomol. 13, 48–52 (1999).

24 Cook, P. E. et al. The use of transcriptional profiles to predict adult mosquito age under field conditions. Proc. Natl. Acad. Sci. U. S. A. 103, 18060–18065 (2006).

25 Sikulu, M. T. et al. Mass spectrometry identification 40. of age-associated proteins from the malaria mosquitoes Anopheles gambiae s.s. and Anopheles stephensi. Data Br. 4, 461–467 (2015).

26 Sinka, M. E. et al. The dominant Anopheles vectors of human malaria in Africa, Europe and the Middle East: occurrence data, distribution maps and bionom-42. ic précis. Parasit. Vectors 4, 89 (2011).

27 Koekemoer, L. L., Kamau, L., Hunt, R. H. & Coetzee, M. A cocktail polymerase chain reaction assay to identify members of the Anopheles funestus (Diptera: 43. Culicidae) group. Am. J. Trop. Med. Hyg. 66, 804–811 (2002).

28 Cohuet, A., Simard, F., Toto, J.C., Kengne, P., Coetzee, M. and Fontennile, D. Species identification within the Anopheles funestus group of malaria vec- 44. tors in Cameroun and evidence fro new species. Am. J. Trop. Med. Hyg. 69, 200–205 (2003).

29 Santolamazza, F. et al. Insertion polymorphisms of SINE200 retrotransposons within speciation islands of Anopheles gambiae molecular forms. Malar. J. 7, 163 (2008).

30 Braack, L. et al. Biting behaviour of African malaria vectors: 1. where do the main vector species bite on the human body? Parasit. Vectors 8, 76 (2015).

31 Lyimo, I. N. & Ferguson, H. M. Ecological and evolutionary determinants of host species choice in mosquito vectors. Trends Parasitol. 25, 189–196 (2009).

32 Lehmann, T. & Diabate, A. The molecular forms of Anopheles gambiae: A phenotypic perspective. Infect. Genet. Evol. 8, 737–746 (2008).

33 Scott, J. A., Brogdon, W. G. & Collins, F. H. Identification of single specimens of the Anopheles gambiae complex by the polymerase chain reaction. Am. J. Trop. Med. Hyg. 49, 520–529 (1993).

34 Bass, C., Williamson, M. S., Wilding, C. S., Donnelly, M. J. & Field, L. M. Identification of the main malaria vectors in the Anopheles gambiae species complex using a TaqMan real-time PCR assay. Malar J 6, 155 (2007).

35 Coosemans, M., Smits, A. & Roelants, P. Intraspecific isozyme polymorphism of Anopheles gambiae in relation to environment, behavior, and malaria transmission in southwestern Burkina Faso. Am. J. Trop. Med. Hyg. 58, 70–74 (1998).

36 Al Ahmed, A. M., Badjah-Hadj-Ahmed, A. Y., Al Othman, Z. A. & Sallam, M. F. Identification of wild collected mosquito vectors of diseases using gas chromatography-mass spectrometry in Jazan Province, Saudi Arabia. J. Mass Spectrom. 48, 1170–1177 (2013).

37 Mayagaya, V. S. et al. Non-destructive determination of age and species of Anopheles gambiae s.l. using near-infrared spectroscopy. Am. J. Trop. Med. Hyg. 81, 622–630 (2009).

38 Reich, G. Near-infrared spectroscopy and imaging: Basic principles and pharmaceutical applications. Adv. Drug Deliv. Rev. 57, 1109–1143 (2005).

39 Fernandes, J. N. et al. Rapid, noninvasive detection of Zika virus in Aedes aegypti mosquitoes by near-infrared spectroscopy. Sci. Adv. 4, eaat0496 (2018).

40 Krajacich, B. J. et al. Analysis of near infrared spectra for age-grading of wild populations of Anopheles gambiae. Parasit. Vectors 10, 552 (2017).

41 Lambert, B. et al. Monitoring the Age of Mosquito Populations Using Near-Infrared Spectroscopy. Sci. Rep. 8, 1–9 (2018).

42 Lin-Vien, D., Colthup, N., Fateley, W. & Grasselli, J. The Handbook of Infrared and Raman Characteristic Frequencies of Organic Molecules. (Academic Press Inc., 1991).

43 Khoshmanesh, A. et al. Screening of *Wolbachia* Endosymbiont Infection in *Aedes aegypti* Mosquitoes Using Attenuated Total Reflection Mid-Infrared Spectroscopy. Anal. Chem. acs.analchem.6b04827 (2017). doi:10.1021/acs.analchem.6b04827.

44 Webster, G. T. et al. Discriminating the intraerythrocytic lifecycle stages of the malaria parasite using synchrotron FT-IR microspectroscopy and an artificial neural network. Anal. Chem. 81, 2516–2524 (2009).

45 Peiris, K., Drolet, B., Cohnstaedt, L. & Dowell, F. Infrared Absorption Characteristics of Culicoides sonorensis in Relation to Insect Age. Am. J. Agric. Sci. Technol. 2, 49–61 (2014).

46 Hastie, T., Tibshirani, R. & Friedman, J. The Elements of Statistical Learning: Data Mining, Inference, and Prediction. (Springer, 2009).

47 Babayan, S. A., Sinclair, A., Duprez, J. S. & Selman, C. Chronic helminth infection burden differentially affects haematopoietic cell development while ageing selectively impairs adaptive responses to infection. Sci. Rep. 8, 1–12 (2018).

48 Babayan, S. A. et al. The immune and non-immune pathways that drive chronic gastrointestinal helminth burdens in the wild. Front. Immunol. 9, (2018).

49 Borchers, M. R. et al. Machine-learning-based calving prediction from activity, lying, and ruminating behaviors in dairy cattle. J. Dairy Sci. 100, 5664–5674 (2017).

50 Sikulu-Lord, M. T. et al. Near-Infrared Spectroscopy, a Rapid Method for Predicting the Age of Male and Female Wild-Type and Wolbachia Infected Aedes aegypti. PLoS Negl Trop Dis 10, e005040 (2016).

51 Sikulu, M. et al. Near-infrared spectroscopy as a complementary age grading and species identification tool for African malaria vectors. Parasit. Vectors 3, 49 (2010).

52 Yahouedo, G. A. et al. Contributions of cuticle permeability and enzyme detoxification to pyrethroid resistance in the major malaria vector Anopheles gambiae. Sci. Rep. 7, 1–10 (2017).

53 Suarez, E. et al. Matrix-assisted laser desorption/ionization-mass spectrometry of cuticular lipid profiles can differentiate sex, age, and mating status of Anopheles gambiae mosquitoes. Anal. Chim. Acta 706, 157–163 (2011).

54 Caputo, B. et al. Identification and composition of cuticular hydrocarbons of the major Afrotropical malaria vector Anopheles gambiae s.s. (Diptera: Culicidae): Analysis of sexual dimorphism and age– related changes. J. Mass Spectrom. 40, 1595–1604 (2005).

55 Molineaux, L. & Gramiccia, G. *The Garki Project. Research on the epidemiology and* control of malaria in the Sudan savanna of West Africa. (World Health Organization, 1980).

56 Lecun, Y., Bengio, Y. & Hinton, G. Deep learning. Nature 521, 436–444 (2015).

57 Balabanidou, V. et al. Cytochrome P450 associated with insecticide resistance catalyzes cuticular hydrocarbon production in Anopheles gambiae. Proc. Natl. Acad. Sci. 113, 9268–9273 (2016).

